# Cardiovascular Disease Related Proteomic Biomarkers of Alcohol Consumption

**DOI:** 10.1101/2020.10.17.332197

**Authors:** Xianbang Sun, Jennifer E. Ho, He Gao, Evangelos Evangelou, Chen Yao, Tianxiao Huan, Shih-Jen Hwang, Paul Courchesne, Martin G. Larson, Daniel Levy, Jiantao Ma, Chunyu Liu

## Abstract

The relationship between alcohol consumption, circulating proteins, and cardiovascular disease (CVD) risk has not been well studied. We performed association analyses of alcohol consumption with three CVD risk factors and 71 CVD-related circulating proteins measured in 6,745 Framingham Heart Study participants (mean age, 49 years; 53% women). We found that an increase in alcohol consumption was associated with a higher risk of incident hypertension (P=7.2E-3) but a lower risk of incident obesity (P=5.7E-4) and type 2 diabetes (P=1.4E-5) in a 14-year of follow-up. Using independent discovery (n=4,348) and validation (n=2,397) samples, we identified 20 alcohol-associated proteins (FDR<0.05 in discovery and P<0.05/n in validation), with majority (18 of 20 proteins) inversely associated with alcohol consumption. The alcohol-protein associations remained similar after removing heavy drinkers. Four proteins demonstrated consistent triangular relationships, as expected, with alcohol consumption and CVD risk factors. For example, a greater level of APOA1, which was associated with a higher alcohol consumption (P=1.2E-65), was associated with a lower risk of type 2 diabetes (P=3.1E-5). However, several others showed inconsistent triangular relationships, e.g., a greater level of GDF15, which was associated with a lower alcohol consumption (P=1.0E-13), was associated with an increased risk of hypertension (P=2.4E-4). In conclusion, we identified 20 alcohol-associated proteins and demonstrated complex relationships between alcohol consumption, circulating proteins and CVD risk factors. Future studies with integration of more proteomic markers and larger sample size are warranted to unravel the complex relationship between alcohol consumption and CVD risk.

## Introduction

Alcohol consumption may have a complex relationship to long-term risk for a variety of diseases. The excessive alcohol use contributes to three million deaths every year globally and is a leading risk factor for more than 60 acute and chronic diseases.^1; 2^ Previous studies indicated that moderate alcohol drinking can have a protective effect on risk for coronary heart disease and all-cause mortality^3;4^ Several recent studies, however, have challenged this view, based in part on results from Mendelian randomization analyses^5; 6^ and large-sample meta-analyses.^2; 5–7^ These recent studies revealed that moderate alcohol intake increases, or at least does not decrease, the risk for cardiovascular disease (CVD).^2; 5–7^

Proteins are composed of one or more long chains of amino acids that are encoded by triplets within the DNA sequence. Circulating proteins play a variety of roles in many physiological changes and responses.^8^ For example, some circulating proteins are receptors or regulators in cell signaling pathways, and many others are transporters of various types of molecules that are critical for a wide range of system functions. As part of the National Heart, Lung, and Blood Institute’s Systems Approach to Biomarker Research in Cardiovascular Disease (SABRe CVD) Initiative, 85 candidate protein biomarkers were measurement in over 7,000 Framingham Heart Study (FHS) participants.^9^ We recently reported that many of these proteins are associated with CVD and all-cause mortality.^9^

We hypothesized that circulating proteins are involved in biological processes that link alcohol consumption to CVD risk owing to their broad range of roles. A better understanding of the relations between alcohol consumption and these CVD-related protein biomarkers will help unravel the complex relationship of alcohol consumption to CVD. To that end, we performed cross-sectional association analyses between circulating proteins and alcohol consumption. We then performed longitudinal association analyses between alcohol-associated proteins and incident risk factors for CVD, including obesity, hypertension, and type 2 diabetes. We also used Mendelian randomization analysis to explore causal relationships between alcohol consumption and alcohol-associated proteins.

## Methods

### Study sample

The FHS is a community‐based, longitudinal cohort study initiated in 1948 to investigate the risk factors for CVD.^10^ The Offspring cohort^11^ was recruited in 1971-1974 and the Third Generation cohort^12^ was recruited in 2002-2005. Since recruitment, both cohorts have been followed for health examinations every 4-8 years to collect demographic and socioeconomic variables and risk factors for CVD. This study was composed of Offspring cohort participants who attended the 7^th^ examination (1998-2001) and Third Generation cohort participants who attended the 1^st^ examination (2002-2005). We excluded participants without alcohol consumption, protein data, or covariates including age, sex, body mass index (BMI), smoking, and cohort index, leaving 2,998 Offspring and 3,747 Third Generation participants for subsequent analyses. The entire study sample was split into a discovery set (n=4,348) and validation set (n=2,397) by independent pedigrees (Table 1). The study design is summarized in Figure 1. All study participants provided written informed consent. All study protocols were approved by the institutional review board of the Boston University Medical Center.

**Table 1.** Participant characteristics

| **Variable** | **Total Sample** | **Discovery Set** | **Replication set** |
| --- | --- | --- | --- |
| N | 6,745 | 4,348 | 2,397 |
|  | **Mean (SD) or n (%)** |  |  |
| Age | 49(14) | 49(14) | 50(14) |
| Women | 3,587(53) | 2,316(53) | 1,271(53) |
| Body mass index (kg/m<sup>2</sup>) | 27(5) | 27.29(5) | 27.66(6) |
| Obesity | 1,726(26) | 1,070(25) | 656(27) |
| Systolic blood pressure (mm Hg) | 121(17) | 121(17) | 122(17) |
| Diastolic blood pressure (mm Hg) | 75(10) | 75(10) | 75(10) |
| Hypertension | 1,976(29) | 1,252(29) | 724(30) |
| Blood pressure medication | 1,312(19) | 816(19) | 496(21) |
| Fasting Serum glucose (mg/dL) | 99(23) | 99(22) | 100(24) |
| Type 2 Diabetes | 443(7) | 256(6) | 187(8) |
| Diabetes medication | 270(4) | 150(3) | 120(5) |
| Current smoking | 988(15) | 621(14) | 367(15) |
| CVD | 423(6) | 253(6) | 170(7) |
|  | **Median (IQR) or n (%)** |  |  |
| Daily alcohol consumption(g/d) | 4(15) | 4(15) | 4.13(15) |
| Non-drinkers | 2672(40) | 1681(39) | 991(41) |

CVD, indicates cardiovascular disease; participant characteristics were contemporaneous with protein measurements.

**Figure 1.**
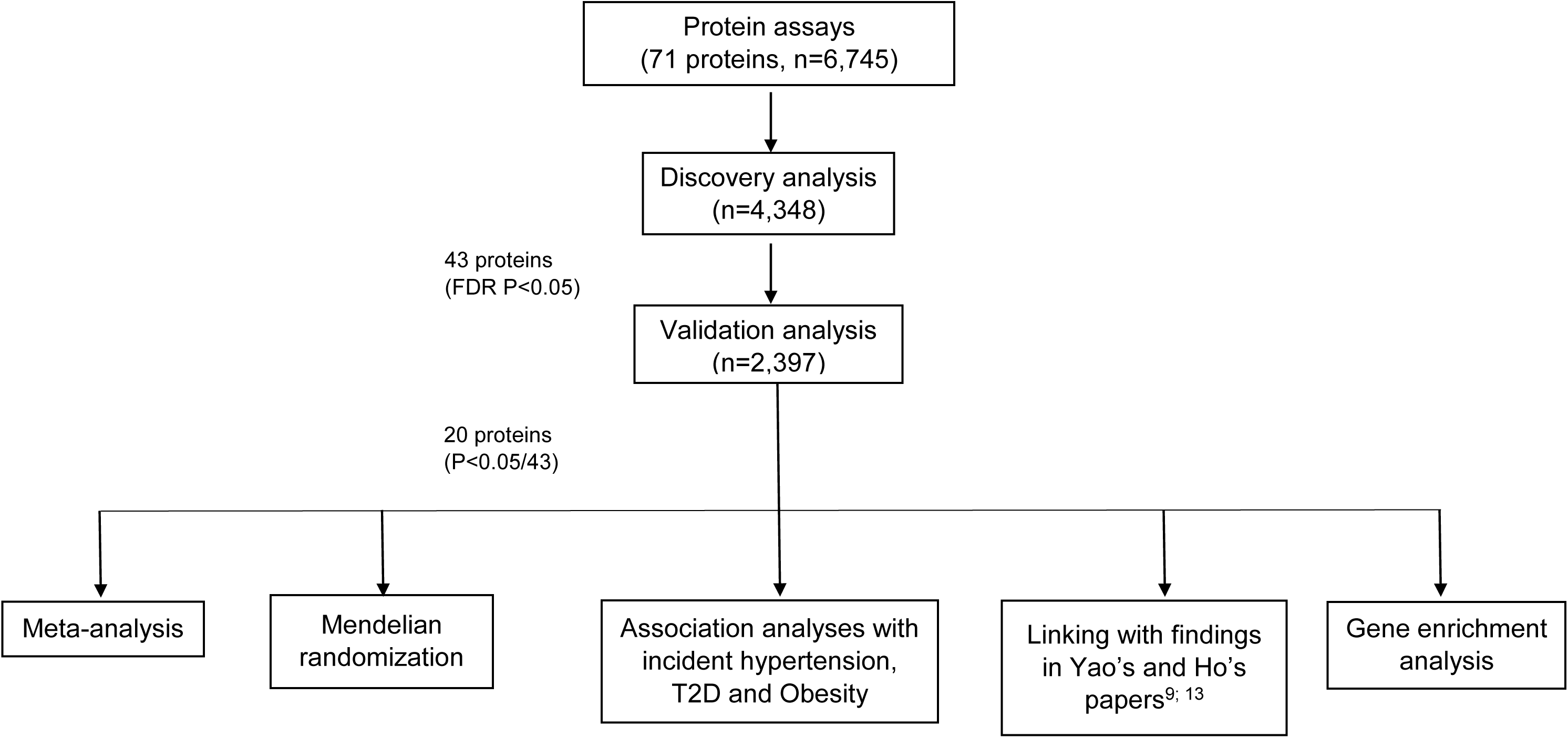
The flow chart of study design Meta-analysis was used to better quantify the alcohol-protein relations; Mendelian randomization was to infer the causal relations of alcohol consumption on protein markers.

### Protein quantification

The selection, quantification, and quality control of the 85 proteins were previously described in detail.^9; 13^ In brief, proteins were selected a priori based on evidence of association with atherosclerotic CVD or its risk factors using comprehensive literature search, proteomics discovery, targeting proteins coded by genes identified via gene expression profiling studies, or GWAS of atherosclerotic CVD and its risk factors. The 85 plasma proteins were assayed using a modified enzyme-linked immunosorbent assay sandwich method, multiplexed on a Luminex xMAP platform (Luminex, Inc., Austin, TX). After calibration and quality control procedures, 71 out of 85 displayed detectable levels for >95% of participant samples, ^9; 13^ and these 71 proteins were used in the present statistical analyses. The 71 proteins had mean inter-assay coefficient of variation of 8.9±5.0% and mean intra‐assay coefficient of variation 7.8±4.9%.^9^

### Alcohol consumption

Alcohol consumption was measured contemporaneously with protein levels. The continuous phenotype, “grams of alcohol consumed per day”, was calculated by using the following converters between drinks and grams: one beer (12 oz) was approximately 14 grams ethanol, one glass of wine (5 oz) was approximately 14 grams ethanol, or one drink of spirit (1.5 oz 80 proof alcohol) was approximately 14 grams ethanol.^14^ To correct right skewness of alcohol drinking data, we added 1 to alcohol consumption data and applied log10 transformation. Additionally, we created a four-category alcohol intake trait: non-drinkers (g/day=0); moderate drinkers (0.1-28 g/day in men and 0.1-14 g/day in women); at-risk drinkers (28.1-42 g/day in men and 14.1-28 g/day in women); and heavy drinkers (>42 g/day in men and >28 g/day in women). We used this categorical alcohol consumption variable to further examine whether alcohol consumption has a non-additive association with the protein biomarkers.

### Genotyping

Affymetrix 500K mapping array and the 50K supplemental Affymetrix array were used in the genotyping in FHS. MaCH software^15^ was used for imputation in conjunction with the 1000G phase 1 version 3 to generate an imputed set of ~30 million variants.^16^ SNPs with imputation quality ratio R-squared<0.5 or minor allele frequencies (MAFs) <0.01 were removed. About 8 million single nucleotide polymorphisms (SNPs) were remained for the present analyses.

### Association and meta-analysis of alcohol consumption with protein markers

The values of concentrations for the 71 proteins were inverse-rank normalized. Each of the transformed protein markers was used as the outcome and alcohol consumption was the predictor variable, adjusting for age, sex, BMI, current smoking status and cohort index (Offspring vs. Third Generation). Linear mixed models were used to identify associations between proteins and alcohol consumption. Random intercept was used to account for family structure. False discovery rate (FDR) <0.05 was used for significance test to account for multiple testing in the discovery stage, i.e., proteins associated with alcohol consumption at FDR <0.05 in the discovery set were brought forward for association analysis in the validation set. We considered P<0.05/n significant in the validation stage, where n is the number of significant proteins in the discovery set.

To better quantify the alcohol-protein relations, meta-analysis was performed with fixed-effect (if heterogeneity *I*^2^ < 0.5) or random-effect (*I*^2^ ≥ 0.5) model to combine summary statistics from discovery and validation sets. We further excluded 351 (5.2%) heavy drinkers (≥3 drinks per day in men and ≥2 drinks per day in women) in the pooled samples (n=6,745) and performed sensitivity analysis to investigate whether alcohol-protein associations were driven by heavy alcohol drinkers. For protein markers that were significant after validation, we also performed an analysis of covariance to compare the average protein levels in moderate, at-risk, and heavy drinkers compared to non-drinkers using linear mixed models, adjusting for the same covariates.

### Longitudinal Association analysis of alcohol-associated proteins with incident CVD risk factors

We performed longitudinal association analysis to examine whether the alcohol-associated proteins obtained from the two-stage association analysis were predictive for three CVD risk factors, including obesity, hypertension and type 2 diabetes in pooled samples. Obesity was defined by BMI ≥30 kg/m^2^, hypertension was defined by systolic blood pressure (SBP) ≥140 mm Hg or diastolic blood pressure (DBP) ≥90 mm Hg or taking hypotensive drugs, and type 2 diabetes was defined by fasting blood glucose level≥126 mg/dL or taking antidiabetic drugs. Prevalent cases at baseline were excluded from the longitudinal analyses for each of the three CVD risk factors, e.g., participants with baseline obesity were excluded in analysis for incident obesity. If a disease was observed, the follow-up time was calculated by the difference between the age of the exam at which the disease was observed for the first time and the age at baseline, otherwise, the follow-up time was calculated as the difference between age at the last follow-up examination and at baseline. Mixed Cox proportional hazards regression models were used to analyze the longitudinal relations. Covariates included age, sex, smoking status and cohort index. Additionally, baseline binary BMI category (<25 or ≥25 kg/m^2^) was adjusted for incident obesity; baseline binary blood pressure (DBP≥80 mm Hg or SBP≥120 mm Hg, otherwise) was adjusted for incident hypertension; baseline binary blood glucose (<100 or ≥100 mg/dL) was adjusted for incident type 2 diabetes. FDR <0.05 was used for significance to account for multiple testing.

### Mendelian randomization analysis to infer the causal relations between alcohol consumption and alcohol-associated proteins

Two-sample Mendelian randomization (MR) analysis was performed to explore the causal relationship of alcohol consumption to the alcohol-related proteins obtained from the two-stage (i.e., discovery and validation) association analysis. We identified 76 SNPs that were associated with alcohol consumption from literature review of candidate gene and genome-wide association analysis (GWAS) of alcohol consumption.^17–21^ Sixty-one of the 76 reported SNPs were significant (P<5×E-8) in the largest and most recent meta-analysis (n~480,000) combining the UK Biobank (UKB), the Cohorts for Heart and Aging Research in Genomic Epidemiology (CHARGE) and Alcohol Genome-Wide Association (AlcGen) Consortium.^18^ The 76 SNPs included those with well-known biological function relevant to alcohol metabolisms (e.g., rs1229984 at *ADH1B*). All MR analyses were conducted using TwoSampleMR R package.^22^ MR-egger test was used to test for horizontal pleiotropy. Inverse-variance weight (IVW) method^23^ was applied to estimate the causal effect. Causal relations were identified by P<0.05.

### Gene enrichment analysis

We performed gene set enrichment analysis using the web-based Gene Ontology tool (http://geneontology.org/) to identify biological pathways for the alcohol associated proteins identified from the two-stage association analysis.

## Results

### Participant characteristics

Most participant characteristics were similar between the discovery set (n=4348; mean age 49 years; 53% women) and validation set (n=2397; mean age 49 years; 53% women) (Table 1). A slightly lower proportion of non-drinkers was included in the discovery set (38.7%) than the validation set (41.3%) (Chi-squared P=0.03). However, the mean daily alcohol consumption was similar between the two sets (10.5 g/day in the discovery set and 10.4 g/day in the validation set, Mann-Whitney U Test, P=0.16). In addition, the means of the inverse-rank normalized proteins were also similar for the two sets (P>0.05/71) except for CXCL16 (−0.04 in the discovery set and 0.08 in the validation set, two-sample t test, P=3.17E-06<0.05/71).

### Increased alcohol assumption was associated with decreased levels for most alcohol-associated proteins

Most (88.29%) of the 71 protein markers displayed modest pairwise correlations (correlation coefficients *r* between −0.2 to 0.2). About two percent of the pairwise correlations were moderate (|*r|*>0.4). The strongest pairwise correlation coefficient (*r* =0.88) was observed between cadherin (CDH13) and low-density lipoprotein receptor (LDLR).^9^

We identified 43 proteins associated with alcohol consumption (FDR<0.05) in the discovery analysis, and 20 of them remained significant (*P*<0.05/43) in the validation analysis (Table 2). The strongest association with alcohol consumption was for apolipoprotein A1 (APOA1) (beta=0.35 with P=2.15×E-41 in discovery set; beta=0.33 with *P*=9.14×E-23 in validation set). Among the 20 protein markers, 18 displayed negative association with alcohol intake, that is, increased alcohol consumption was associated with decreased circulating protein levels in plasma. Two protein markers displayed positive association with alcohol consumption. The directionality of the alcohol-protein associations was same between the discovery and the validation sets for these 20 alcohol-associated proteins (Table 2). Magnitude of the associations was similar for most of protein markers (i.e., *I*^2^>0.5 in the meta-analysis that combined summary statistics of discovery and validation sets for the alcohol-related proteins; **Supplemental Table 1**).

**Table 2.** Significant alcohol-associated protein markers (Bonferroni corrected *P*<0.001 in validation)

| Protein | Discovery analysis (n=4348) |  |  |  | Validation analysis (n=2397) |  |  |
| --- | --- | --- | --- | --- | --- | --- | --- |
|  | Beta | SE | p | FDR | Beta | SE | p |
| APOA1 | 0.352 | 0.026 | 2.2E-41 | 1.5E-39 | 0.335 | 0.032 | 2.1E-24 |
| RESISTIN | -0.230 | 0.026 | 2.8E-19 | 1.0E-17 | -0.312 | 0.033 | 3.3E-20 |
| MPO | -0.195 | 0.025 | 8.4E-15 | 8.5E-14 | -0.248 | 0.033 | 1.8E-13 |
| EFEMP1 | -0.149 | 0.026 | 9.4E-09 | 5.5E-08 | -0.217 | 0.033 | 1.1E-10 |
| ANGPTL3 | -0.210 | 0.024 | 1.2E-17 | 2.2E-16 | -0.209 | 0.032 | 1.2E-10 |
| CD56 | -0.193 | 0.025 | 9.6E-15 | 8.5E-14 | -0.222 | 0.034 | 1.3E-10 |
| sRAGE | -0.226 | 0.025 | 4.3E-19 | 1.0E-17 | -0.213 | 0.034 | 2.8E-10 |
| FBN | -0.166 | 0.025 | 6.3E-11 | 4.4E-10 | -0.210 | 0.033 | 2.9E-10 |
| Notch1 | -0.210 | 0.026 | 1.1E-15 | 1.5E-14 | -0.215 | 0.034 | 4.3E-10 |
| GDF15 | -0.130 | 0.025 | 2.4E-07 | 1.0E-06 | -0.184 | 0.033 | 4.6E-08 |
| OSTEOCALCIN | -0.137 | 0.026 | 1.1E-07 | 5.0E-07 | -0.183 | 0.034 | 8.2E-08 |
| B2M | -0.122 | 0.026 | 4.0E-06 | 1.5E-05 | -0.162 | 0.034 | 1.6E-06 |
| MMP8 | -0.058 | 0.026 | 2.4E-02 | 4.1E-02 | -0.157 | 0.034 | 5.9E-06 |
| Cystatin-C | -0.157 | 0.026 | 1.2E-09 | 7.5E-09 | -0.155 | 0.034 | 6.0E-06 |
| TIMP1 | -0.137 | 0.026 | 1.2E-07 | 5.3E-07 | -0.154 | 0.034 | 8.1E-06 |
| MYOGLOBIN | -0.148 | 0.026 | 1.1E-08 | 6.1E-08 | -0.142 | 0.035 | 4.0E-05 |
| PAI1 | 0.067 | 0.024 | 5.5E-03 | 1.2E-02 | 0.127 | 0.03125 | 4.94E-05 |
| FGF23 | -0.089 | 0.026 | 8.0E-04 | 2.5E-03 | -0.126 | 0.034 | 1.9E-04 |
| CNTN1 | -0.199 | 0.025 | 3.5E-15 | 4.1E-14 | -0.117 | 0.033 | 4.4E-04 |
| Hemopexin | -0.0556 | 0.025 | 0.0289 | 0.0478 | -0.113 | 0.033 | 0.00067 |

Linear mixed multivariable-adjusted model, adjusting for age, sex, body mass index, smoking, and cohort index, and family structure. SE, standard error; FDR, false discovery rate.

After excluding heavy drinkers, 19 of these 20 alcohol-associated proteins remained significant, with slight attenuation for the magnitude of the associations with alcohol consumption, indicating that the association was not driven by heavy alcohol consumption (Figure 2, **Supplemental Table 2**). We further compared the average plasma protein levels in moderate, at-risk, and heavy drinkers to that in non-drinkers. Alcohol drinking categories displayed an additive effect on seventeen proteins, but non-linear effect on three proteins (**Supplemental Table 3**). The protein levels of hemopexin were similar in light drinkers and heavy drinkers, which was slightly different from their levels in non-drinkers. The protein levels of GDF15 were similar in moderate and at-risk drinkers, while the level in heavy drinkers was not significantly different from that in non-drinkers. Lowest plasma level was found in at-risk drinkers for FBN (**Supplemental Table 3**).

**Figure 2.**
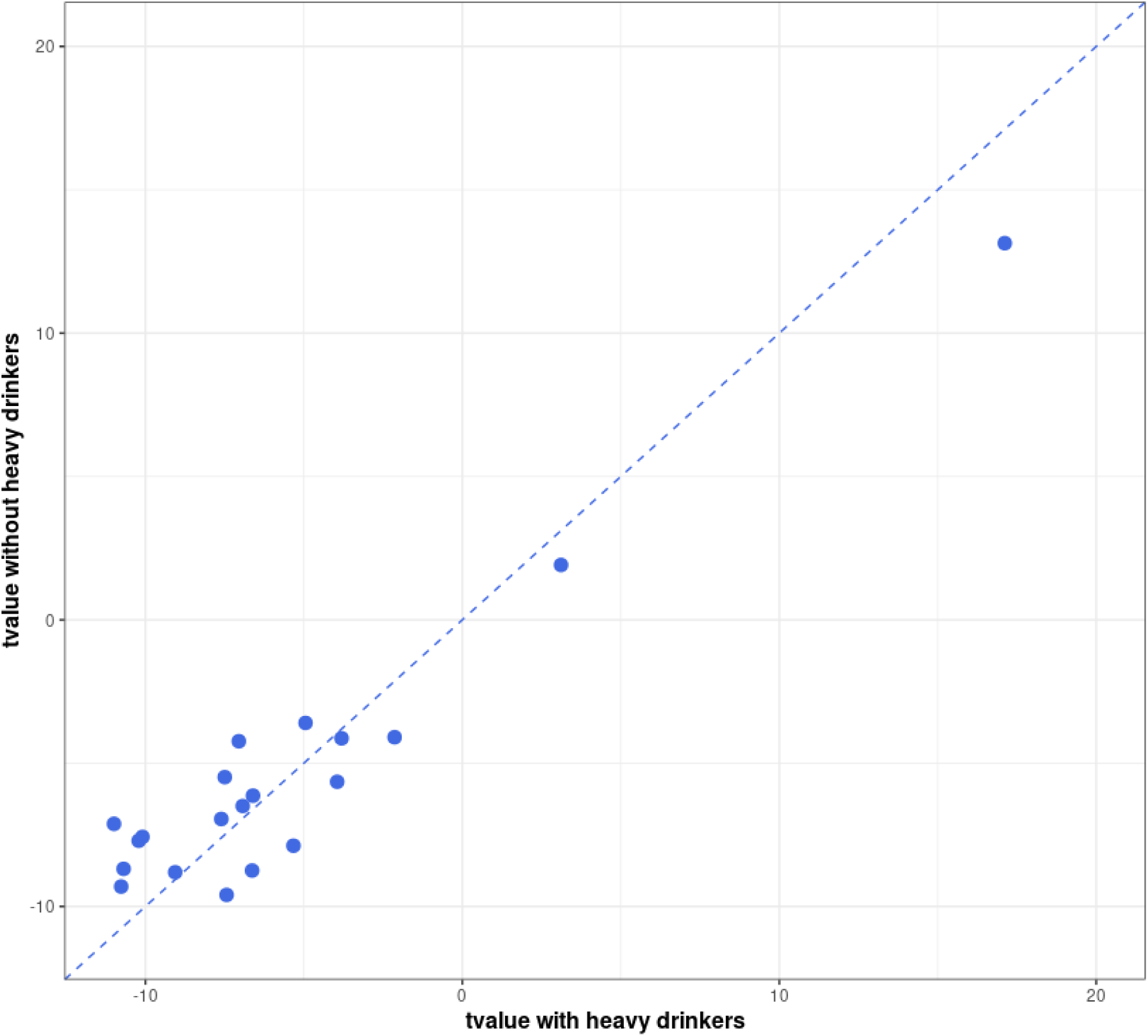
Comparison of t-value in association analysis of 20 significant proteins in participants with (n=6745) and without (n=6249) heavy drinkers

### Gene enrichment analysis

The 20 alcohol-associated proteins were enriched for 24 biological processes (FDR<0.05, **Supplemental Table 5**). The top significant biological processes were ‘negative regulation of cell adhesion molecule production’ (FDR=4.15E-3), ‘post-embryonic eye morphogenesis’ (FDR=7.54E-3), ‘post-embryonic animal organ morphogenesis’ (FDR=1.03E-2), and ‘modulation of age-related behavioral decline’ (FDR=1.04E-2).

### Possible causal effects of alcohol intake on alcohol-associated proteins

We explored causal relations between alcohol consumption and the 20 alcohol-related proteins (from the discovery-replication analysis) by two-sample MR analyses. We found evidence of causal effects of alcohol consumption on two proteins (P<0.05), including angiopoietin-like protein 3 (ANGPTL3, beta=-0.84, P=0.004) and Hemopexin (beta=-0.75, FDR=0.006). The MR analyses showed same directionality as that observed in the association analyses for these two proteins (**Supplemental Table 4**).

### Alcohol-associated proteins were associated with multiple CVD risk factors

After excluding prevalent cases, we had 5,019, 4,769, 6,317 participants who were free of obesity, hypertension, and type 2 diabetes, respectively, at the baseline. In total, 718 (14.39%), 1,257 (26.72%), 400 (6.36%) participants developed obesity, hypertension, and type 2 diabetes, respectively, during a median follow-up of 13 years (interquartile range 7 years). Eight of the 20 alcohol-associated proteins displayed significant associations (FDR<0.05) with the development of one or two of the diseases: six with incident obesity, two with incident hypertension, and three with incident type 2 diabetes (Table 3). In addition, nine of the 20 alcohol-associated proteins were significantly correlated with incidence of CVD.^9^ All of them displayed positive relationship with CVD risk (Figure 4).

**Table 3.** Associations between alcohol-associated protein markers with cardiovascular disease risk factors

| **Trait** | **Biomarker** | **HR** | **95%**<br>**LCL** | **95%**<br>**UCL** | **P Value** | **FDR P** |
| --- | --- | --- | --- | --- | --- | --- |
| Obesity<br>(N=718) | sRAGE | 0.81 | 0.75 | 0.88 | 2.21E-07 | 1.33E-05 |
|  | CNTN1 | 0.85 | 0.79 | 0.92 | 8.57E-05 | 1.71E-03 |
|  | Osteocalcin | 0.88 | 0.81 | 0.95 | 1.84E-03 | 1.38E-02 |
|  | FBN | 1.13 | 1.04 | 1.23 | 2.57E-03 | 1.71E-02 |
|  | APOA1 | 0.89 | 0.83 | 0.97 | 3.93E-03 | 2.21E-02 |
|  | CD56 | 0.89 | 0.83 | 0.96 | 4.05E-03 | 2.21E-02 |
| Hypertension<br>(N=1257) | sRAGE | 0.89 | 0.84 | 0.94 | 1.16E-04 | 1.75E-03 |
|  | GDF15 | 1.12 | 1.06 | 1.19 | 2.44E-04 | 2.44E-03 |
| Type 2<br>Diabetes<br>(N=400) | APOA1 | 0.80 | 0.72 | 0.89 | 3.13E-05 | 9.40E-04 |
|  | GDF15 | 1.23 | 1.10 | 1.37 | 2.27E-04 | 2.44E-03 |
|  | PAI | 1.23 | 1.10 | 1.38 | 5.02E-04 | 4.30E-03 |

Mixed multivariable-adjusted Cox model, adjusted for age, sex, smoking, and cohort index, and family structure. Additionally, baseline binary BMI category (BMI>25) was adjusted for incident obesity; baseline binary blood pressure (DBP≥80 or SBP≥120) was adjusted for incident hypertension; baseline binary blood glucose (≥100) was adjusted for incident type 2 diabetes. CVD, cardiovascular disease; HR, hazards ratio per 1-SD change in rank normalized data of protein markers; LCL, lower 95% confidence interval; UCL, upper 95% confidence interval

In our study samples, higher level of alcohol consumption was associated with higher risk of developing incident hypertension (HR= 1.15, P=7.23E-3, 95% CI=1.04, 1.27) but with lower risk of incident obesity (HR = 0.80, P=5.71E-4, 95% CI=0.70, 0.91) and incident type 2 diabetes (HR = 0.69, P=1.38E-5, 95% CI=0.58, 0.82). Because of the inverse relationship for most of alcohol-protein pairs in our data, we expected that, for most of alcohol-associated proteins, their increased levels were associated with lower incidence for hypertension and higher incidence for obesity and type 2 diabetes. Our three-way association analyses between alcohol consumption, protein markers, and incident CVD risk factors demonstrated several consistent relations with our expectation. For example, we found that higher plasma levels of APOA1, associated with higher alcohol consumption in association analysis, were associated with a reduced risk of developing obesity (HR=0.89, 95%CI=0.83, 0.97; P=0.004) and type 2 diabetes (HR=0.80, 95%CI=0.72, 0.89; P=3.1E-5) (Table 3 and Figure 3). In addition to the consistent three-way relationships of alcohol-APOA1-type 2 diabetes, alcohol-sRAGE-hypertension, alcohol-FBN-obesity, and alcohol-GDF15-diabetes were consistent as expected (**Supplemental Figure 1a**). However, the directionality of several three-way relationships was not consistent to what we expected (**Supplemental Figure 1b**). For example, higher plasma levels of GDF15, associated with lower alcohol consumption, were associated with an increased risk of developing hypertension (HR=1.12, 95%CI=1.06, 1.19; P=2.4E-4) (Table 3 and **Supplemental Figure 1b**).

**Figure 3.**
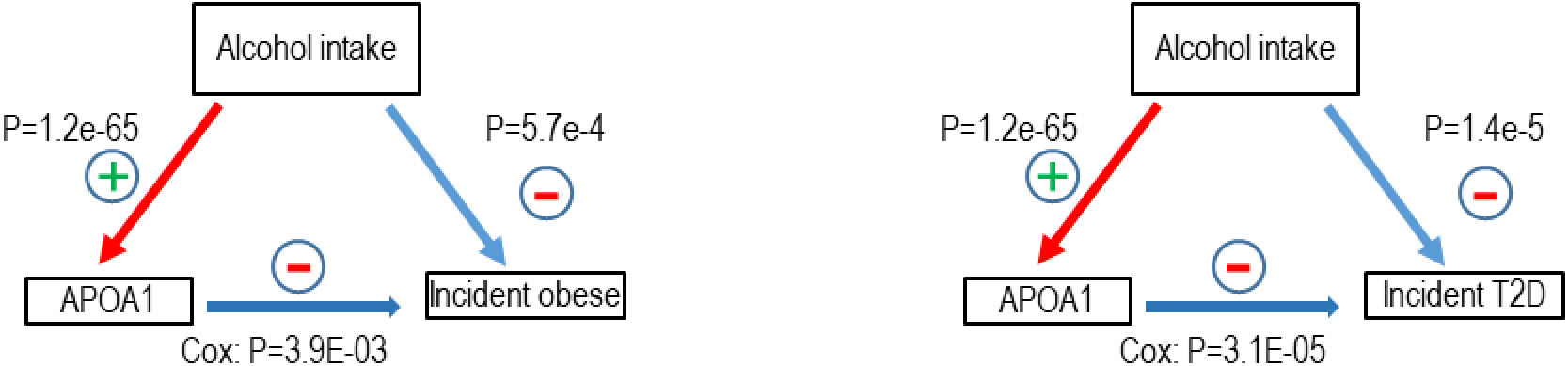
Three–way association of alcohol intake, APOA1, incident obese and T2D

**Figure 4.**
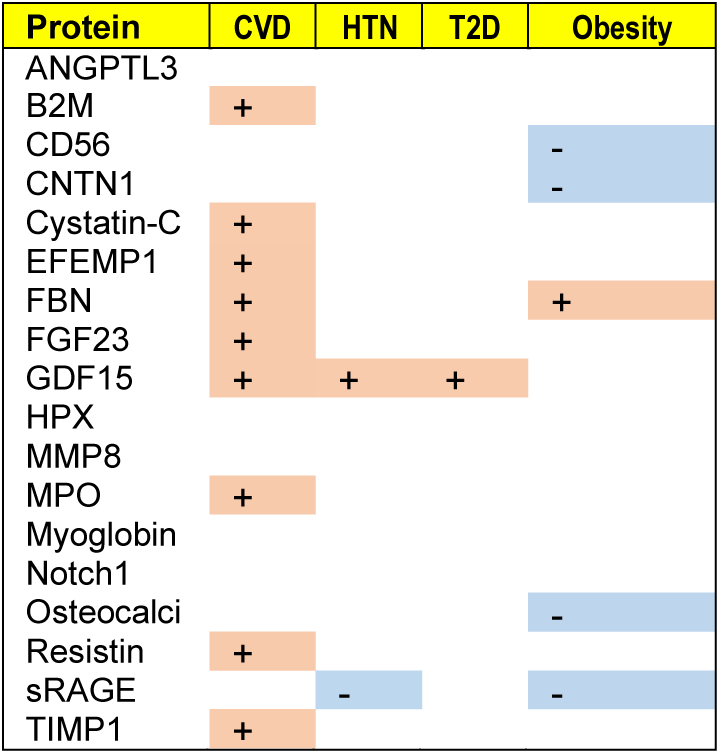
Association of alcohol associated proteins with CVD^9^, and CVD risk factors All proteins in the figure displayed reduced plasma levels with higher level of alcohol consumption. ‘+’ represents a positive association, that is, a higher plasma protein level was associated with a higher risk of CVD and CVD risk factor; ‘-’ represents a negative association, that is, a higher plasma protein level was associated with a lower risk of CVD and CVD risk factor.

## Discussion

In this study, we explored the associations between alcohol consumption and 71 proteins that were preselected due to their potential relations with CVD. As of to date, this is the largest population-based study to explore the associations between circulating protein markers and alcohol consumption. Using discovery and validation analyses, we identified 20 alcohol-protein pairs, with increased alcohol consumption was linearly associated with lower plasma levels for most of these identified protein markers. We also showed strong associations between several alcohol-associated proteins with the three major CVD risk factors, potentially by regulating immune responses such as negative regulation of cell adhesion molecule production.^24^ Taken together, our study provide novel evidence that integrating circulating protein markers may be useful to unravel the complex relations between alcohol consumption and CVD.

We previously investigated the relations of the 71 circulating proteins to CVD outcomes.^13^ A longitudinal analysis showed that eleven alcohol-associated proteins identified in this study were associated with atherosclerotic CVD, heart failure, and/or CVD-specific and all-cause mortality in a follow-up study^9^ (**Supplemental Table 6**). A study utilizing Mendelian randomization analysis showed that two of the 20 alcohol-associated proteins, myeloperoxidase (MPO) and cystatin C were causally associated with coronary heart disease (CHD).^13^ In addition, this study showed that the protein quantitative trait loci (pQTLs) for beta-2 microglobulin (B2M) coincide with the CHD-associated SNPs reported in the GWAS catalog.^13^

Several studies have investigated proteomic markers (up to 246 proteins) of extreme alcohol intake in cerebral cortex and hepatocellular carcinoma cell lines.^25–28^ Statistical power of these studies was, however, limited due to small sample sizes (< 50). In addition, generalizability to general population was limited because extreme alcohol intake was the research focus in these previous studies. ^25–28^ The present study used pre-selected CVD-related protein markers, which may help unravel the relationship of alcohol-associated protein markers with CVD and CVD risk factor. However, using pre-selected proteins limits the capacity to investigate the relationship of a large portion of human proteome with alcohol consumption and with CVD. Future investigations with the entire proteome are warranted to uncover comprehensive proteomic signatures of alcohol consumption and the complex relationships of alcohol consumption and CVD risk.

Many environmental factors such as diet^29^ may influence plasma protein levels. Therefore, we performed MR analyses to explore the possible causal influence of alcohol intake on the 20 alcohol-associated proteins. MR analyses implicated that alcohol consumption may have a causal effect for two proteins, ANGPTL3, and hemopexin. The MR analysis showed similar association as that observed in the association analysis, i.e., higher alcohol consumption associated with lower levels of ANGPTL3 and hemopexin. ANGPTL3 is a secretory protein regulating plasma lipid levels via affecting lipoprotein lipase- and endothelial lipase-mediated hydrolysis of triglycerides and phospholipids.^30^ This study showed that increased alcohol intake may be protective for the development of CVD through the decreased level of ANGPTL3 in plasma, while a previous finding revealed a positive correlation between ANGPTL3 and CVD risk,^31^ Hemopexin is an acute phase glycoprotein with highest binding affinity to heme.^32^ This protein is associated with HDL in the inflammatory process. However, other study suggests that higher hemopexin, a plasma heme scavenger, may be protective to CVD risk.^33^ Therefore, our data revealed a complex relations between alcohol consumption, protein markers, and CVD

The participants in the study were all European descent. Therefore, the finding in this study may not be generalizable to individuals of non-European ancestries. The instrumental variables (IVs) (i.e., alcohol-associated SNPs) in MR analysis to infer causal associations of alcohol on protein markers were weak because the SNPs jointly explained 1.6% of variation in alcohol consumption. In addition, although we tested horizontal pleiotropy, we could not completely rule out its impact in our MR analysis.

In conclusion, we have identified 20 alcohol-associated proteins in middle to older aged adults. These findings demonstrate that integrating protein markers with observational data may help unravel the complex relationship between alcohol consumption and CVD risk. These alcohol-associated proteins may have implications for the prevention and treatment of alcohol-related CVD risk. Additional studies are warranted to replicate our findings, particularly in individuals with different demographic, socioeconomic, or ethnic groups.

## Supporting information

Alcohol_Protein_Suppl_Table_Figure

## The URLs

1. The NHGRI-EBI Catalog of human genome-wide association studies (The GWAS catalog): https://www.ebi.ac.uk/gwas/
2. The GWAS database for type 2 diabetes: https://gwas.mrcieu.ac.uk/datasets/ebi-a-GCST006867/
3. The GWAS database for hypertension: https://gwas.mrcieu.ac.uk/datasets/ukb-b-12493/.
4. The GWAS database for obesity: https://gwas.mrcieu.ac.uk/datasets/ukb-b-15541/.

## Notes

### Competing Interest Statement

The authors have declared no competing interest.

### Summary of Updates

Remove periods from titles of tables and figures

## References

1. Rehm, J., and Imtiaz, S. (2016). A narrative review of alcohol consumption as a risk factor for global burden of disease. Subst Abuse Treat Prev Policy 11, 37.

2. Collaborators, G.B.D.A. (2018). Alcohol use and burden for 195 countries and territories, 1990-2016: a systematic analysis for the Global Burden of Disease Study 2016. Lancet 392, 1015–1035.

3. Di Castelnuovo, A., Costanzo, S., Bagnardi, V., Donati, M.B., Iacoviello, L., and de Gaetano, G. (2006). Alcohol dosing and total mortality in men and women: an updated meta-analysis of 34 prospective studies. Arch Intern Med 166, 2437–2445.

4. Ronksley, P.E., Brien, S.E., Turner, B.J., Mukamal, K.J., and Ghali, W.A. (2011). Association of alcohol consumption with selected cardiovascular disease outcomes: a systematic review and meta-analysis. BMJ 342, d671.

5. Chikritzhs, T.N., Naimi, T.S., Stockwell, T.R., and Liang, W. (2015). Mendelian randomisation meta-analysis sheds doubt on protective associations between ‘moderate’ alcohol consumption and coronary heart disease. Evid Based Med 20, 38.

6. Holmes, M.V., Dale, C.E., Zuccolo, L., Silverwood, R.J., Guo, Y., Ye, Z., Prieto-Merino, D., Dehghan, A., Trompet, S., Wong, A., et al. (2014). Association between alcohol and cardiovascular disease: Mendelian randomisation analysis based on individual participant data. BMJ 349, g4164.

7. Stockwell, T., Zhao, J., Panwar, S., Roemer, A., Naimi, T., and Chikritzhs, T. (2016). Do “Moderate” Drinkers Have Reduced Mortality Risk? A Systematic Review and Meta-Analysis of Alcohol Consumption and All-Cause Mortality. J Stud Alcohol Drugs 77, 185–198.

8. Voet, D., Voet, J.G., and Pratt, C.W. (2006). Fundamentals of biochemistry: life at the molecular level.(Hoboken, N.J.: Wiley).

9. Ho, J.E., Lyass, A., Courchesne, P., Chen, G., Liu, C., Yin, X., Hwang, S.J., Massaro, J.M., Larson, M.G., and Levy, D. (2018). Protein Biomarkers of Cardiovascular Disease and Mortality in the Community. J Am Heart Assoc 7.

10. Dawber, T.R., Meadors, G.F., and Moore, F.E., Jr. (1951). Epidemiological approaches to heart disease: the Framingham Study. Am J Public Health Nations Health 41, 279–281.

11. Feinleib, M., Kannel, W.B., Garrison, R.J., McNamara, P.M., and Castelli, W.P. (1975). The Framingham Offspring Study. Design and preliminary data. Prev Med 4, 518–525.

12. Splansky, G.L., Corey, D., Yang, Q., Atwood, L.D., Cupples, L.A., Benjamin, E.J., D’Agostino, R.B., Sr., Fox, C.S., Larson, M.G., Murabito, J.M., et al. (2007). The Third Generation Cohort of the National Heart, Lung, and Blood Institute’s Framingham Heart Study: design, recruitment, and initial examination. Am J Epidemiol 165, 1328–1335.

13. Yao, C., Chen, G., Song, C., Keefe, J., Mendelson, M., Huan, T., Sun, B.B., Laser, A., Maranville, J.C., Wu, H., et al. (2018). Genome-wide mapping of plasma protein QTLs identifies putatively causal genes and pathways for cardiovascular disease. Nat Commun 9, 3268.

14. Liu, C., Marioni, R.E., Hedman, A.K., Pfeiffer, L., Tsai, P.C., Reynolds, L.M., Just, A.C., Duan, Q., Boer, C.G., Tanaka, T., et al. (2018). A DNA methylation biomarker of alcohol consumption. Mol Psychiatry 23, 422–433.

15. Willer, C.J., Sanna, S., Jackson, A.U., Scuteri, A., Bonnycastle, L.L., Clarke, R., Heath, S.C., Timpson, N.J., Najjar, S.S., Stringham, H.M., et al. (2008). Newly identified loci that influence lipid concentrations and risk of coronary artery disease. Nat Genet 40, 161–169.

16. Genomes Project, C., Auton, A., Brooks, L.D., Durbin, R.M., Garrison, E.P., Kang, H.M., Korbel, J.O., Marchini, J.L., McCarthy, S., McVean, G.A., et al. (2015). A global reference for human genetic variation. Nature 526, 68–74.

17. Bierut, L.J., Goate, A.M., Breslau, N., Johnson, E.O., Bertelsen, S., Fox, L., Agrawal, A., Bucholz, K.K., Grucza, R., Hesselbrock, V., et al. (2012). ADH1B is associated with alcohol dependence and alcohol consumption in populations of European and African ancestry. Mol Psychiatry 17, 445–450.

18. Schumann, G., Liu, C., O’Reilly, P., Gao, H., Song, P., Xu, B., Ruggeri, B., Amin, N., Jia, T., Preis, S., et al. (2016). KLB is associated with alcohol drinking, and its gene product beta-Klotho is necessary for FGF21 regulation of alcohol preference. Proc Natl Acad Sci U S A 113, 14372–14377.

19. Jorgenson, E., Thai, K.K., Hoffmann, T.J., Sakoda, L.C., Kvale, M.N., Banda, Y., Schaefer, C., Risch, N., Mertens, J., Weisner, C., et al. (2017). Genetic contributors to variation in alcohol consumption vary by race/ethnicity in a large multi-ethnic genome-wide association study. Mol Psychiatry 22, 1359–1367.

20. Clarke, T.K., Adams, M.J., Davies, G., Howard, D.M., Hall, L.S., Padmanabhan, S., Murray, A.D., Smith, B.H., Campbell, A., Hayward, C., et al. (2017). Genome-wide association study of alcohol consumption and genetic overlap with other health-related traits in UK Biobank (N=112 117). Mol Psychiatry 22, 1376–1384.

21. Evangelou, E., Gao, H., Chu, C., Ntritsos, G., Blakeley, P., Butts, A.R., Pazoki, R., Suzuki, H., Koskeridis, F., Yiorkas, A.M., et al. (2019). New alcohol-related genes suggest shared genetic mechanisms with neuropsychiatric disorders. Nat Hum Behav 3, 950–961.

22. Hemani, G., Zheng, J., Elsworth, B., Wade, K.H., Haberland, V., Baird, D., Laurin, C., Burgess, S., Bowden, J., Langdon, R., et al. (2018). The MR-Base platform supports systematic causal inference across the human phenome. Elife 7.

23. Burgess, S., Butterworth, A., and Thompson, S.G. (2013). Mendelian randomization analysis with multiple genetic variants using summarized data. Genet Epidemiol 37, 658–665.

24. Harjunpaa, H., Llort Asens, M., Guenther, C., and Fagerholm, S.C. (2019). Cell Adhesion Molecules and Their Roles and Regulation in the Immune and Tumor Microenvironment. Front Immunol 10, 1078.

25. Abdul Muneer, P.M., Alikunju, S., Szlachetka, A.M., and Haorah, J. (2012). The mechanisms of cerebral vascular dysfunction and neuroinflammation by MMP-mediated degradation of VEGFR-2 in alcohol ingestion. Arterioscler Thromb Vasc Biol 32, 1167–1177.

26. Kashem, M.A., Sultana, N., Pow, D.V., and Balcar, V.J. (2019). GLAST (GLutamate and ASpartate Transporter) in human prefrontal cortex; interactome in healthy brains and the expression of GLAST in brains of chronic alcoholics. Neurochem Int 125, 111–116.

27. Yan, G., Lestari, R., Long, B., Fan, Q., Wang, Z., Guo, X., Yu, J., Hu, J., Yang, X., Chen, C., et al. (2016). Comparative Proteomics Analysis Reveals L-Arginine Activates Ethanol Degradation Pathways in HepG2 Cells. Sci Rep 6, 23340.

28. Lai, X., Liangpunsakul, S., Li, K., and Witzmann, F.A. (2015). Proteomic profiling of human sera for discovery of potential biomarkers to monitor abstinence from alcohol abuse. Electrophoresis 36, 556–563.

29. Anderson, N.L., and Anderson, N.G. (2002). The human plasma proteome: history, character, and diagnostic prospects. Mol Cell Proteomics 1, 845–867.

30. Tikka, A., and Jauhiainen, M. (2016). The role of ANGPTL3 in controlling lipoprotein metabolism. Endocrine 52, 187–193.

31. Dewey, F.E., Gusarova, V., Dunbar, R.L., O’Dushlaine, C., Schurmann, C., Gottesman, O., McCarthy, S., Van Hout, C.V., Bruse, S., Dansky, H.M., et al. (2017). Genetic and Pharmacologic Inactivation of ANGPTL3 and Cardiovascular Disease. N Engl J Med 377, 211–221.

32. Tolosano, E., and Altruda, F. (2002). Hemopexin: structure, function, and regulation. DNA Cell Biol 21, 297–306.

33. Vinchi, F., De Franceschi, L., Ghigo, A., Townes, T., Cimino, J., Silengo, L., Hirsch, E., Altruda, F., and Tolosano, E. (2013). Hemopexin therapy improves cardiovascular function by preventing heme-induced endothelial toxicity in mouse models of hemolytic diseases. Circulation 127, 1317–1329.

