## Supplementary material for "Cardiovascular Disease Related Proteomic Biomarkers of Alcohol Consumption": Alcohol_Protein_Suppl_Table_Figure

Supplemental Tables and Figures

**Supplemental Table 1**. Association results for the 20 alcohol-related proteins with alcohol intake from meta-analysis, discovery and validation stages

| | Meta-analysis | | | Discovery analysis (n=4348) | | | Validation analysis (n=2397) | | |
|---|---|---|---|---|---|---|---|---|---|
| Protein | Beta | SE | p | Beta | SE | p | Beta | SE | p |
| APOA1 | 0.345 | 0.0202 | 1.18E-65 | 0.352 | 0.0258 | 2.15E-41 | 0.335 | 0.0324 | 2.13E-24 |
| sRAGE | -0.221 | 0.0201 | 4.32E-28 | -0.226 | 0.0251 | 4.31E-19 | -0.213 | 0.0336 | 2.81E-10 |
| ANGPTL3 | -0.21 | 0.0195 | 5.17E-27 | -0.21 | 0.0245 | 1.23E-17 | -0.209 | 0.0323 | 1.17E-10 |
| MPO | -0.214 | 0.02 | 1.20E-26 | -0.195 | 0.025 | 8.41E-15 | -0.248 | 0.0334 | 1.77E-13 |
| NOTCH1 | -0.211 | 0.0207 | 1.80E-24 | -0.21 | 0.026 | 1.07E-15 | -0.215 | 0.0342 | 4.31E-10 |
| CD56 | -0.203 | 0.0201 | 6.13E-24 | -0.193 | 0.0248 | 9.55E-15 | -0.222 | 0.0343 | 1.29E-10 |
| FBN | -0.182 | 0.0201 | 1.32E-19 | -0.166 | 0.0253 | 6.26E-11 | -0.21 | 0.0331 | 2.89E-10 |
| Cystatin-C | -0.156 | 0.0205 | 2.87E-14 | -0.157 | 0.0257 | 1.17E-09 | -0.155 | 0.0341 | 6.01E-06 |
| Osteocalcin | -0.153 | 0.0205 | 6.64E-14 | -0.137 | 0.0256 | 1.06E-07 | -0.183 | 0.034 | 8.19E-08 |
| GDF15 | -0.15 | 0.0201 | 1.03E-13 | -0.13 | 0.0252 | 2.38E-07 | -0.184 | 0.0334 | 4.57E-08 |
| Myoglobin | -0.146 | 0.0207 | 1.78E-12 | -0.148 | 0.0258 | 1.11E-08 | -0.142 | 0.0345 | 4.01E-05 |
| TIMP1 | -0.143 | 0.0207 | 4.25E-12 | -0.137 | 0.0258 | 1.19E-07 | -0.154 | 0.0345 | 8.10E-06 |
| *Resistin | -0.268 | 0.0404 | 3.33E-11 | -0.23 | 0.0255 | 2.80E-19 | -0.312 | 0.0335 | 3.33E-20 |
| B2M | -0.137 | 0.0208 | 4.00E-11 | -0.122 | 0.0263 | 3.95E-06 | -0.162 | 0.0337 | 1.62E-06 |
| *EFEMP1 | -0.179 | 0.0337 | 9.95E-08 | -0.149 | 0.0259 | 9.36E-09 | -0.217 | 0.0334 | 1.11E-10 |
| FGF23 | -0.103 | 0.0208 | 7.57E-07 | -0.089 | 0.0265 | 0.0008 | -0.126 | 0.0337 | 0.0002 |
| *CNTN1 | -0.161 | 0.0408 | 7.84E-05 | -0.199 | 0.0252 | <1E-16 | -0.117 | 0.0333 | 0.0004 |
| Hemopexin | -0.077 | 0.0202 | 0.00014 | -0.056 | 0.0254 | 0.0289 | -0.113 | 0.0331 | 0.0007 |
| *PAI1 | 0.094 | 0.03 | 0.00185 | 0.067 | 0.024 | 0.0055 | 0.127 | 0.0313 | <1E-16 |
| *MMP8 | -0.105 | 0.049 | 0.03259 | -0.058 | 0.0258 | 0.0241 | -0.157 | 0.0345 | <1E-16 |

Discovery and validation analyses: linear Mixed multivariable-adjusted model, adjusted for age, sex, body mass index, smoking, and cohort index, random intercept adjusted for family structure. Meta-analysis: fixed-effect (if heterogeneity $I^2 < 0.5$) or random-effect ($I^2 \geq 0.5$) model to combine summary statistics from discovery and validation sets. * indicates $I^2 \geq 0.5$.

**Supplemental Table 2.** Comparison of the associations between top 20 proteins and alcohol consumption in all participants and in participants excluding heavy drinkers

| | All participants (n=6745) | | | Participants without heavy drinkers (n= 6249) | | |
|---|---|---|---|---|---|---|
| Protein | Beta | SE | p | Beta | SE | p |
| APOA1 | 0.345 | 0.0202 | 1.18E-65 | 0.298 | 0.0227 | 8.58E-39 |
| sRAGE | -0.221 | 0.0201 | 4.32E-28 | -0.161 | 0.0226 | 1.26E-12 |
| ANGPTL3 | -0.21 | 0.0195 | 5.17E-27 | -0.204 | 0.0220 | 1.97E-20 |
| MPO | -0.214 | 0.02 | 1.20E-26 | -0.196 | 0.0226 | 5.01E-18 |
| NOTCH1 | -0.211 | 0.0207 | 1.80E-24 | -0.181 | 0.0234 | 1.57E-14 |
| CD56 | -0.203 | 0.0201 | 6.13E-24 | -0.171 | 0.0226 | 4.38E-14 |
| FBN | -0.182 | 0.0201 | 1.32E-19 | -0.200 | 0.0227 | 1.77E-18 |
| Cystatin-C | -0.156 | 0.0205 | 2.87E-14 | -0.161 | 0.0232 | 4.20E-12 |
| Osteocalcin | -0.153 | 0.0205 | 6.64E-14 | -0.127 | 0.0232 | 4.28E-08 |
| GDF15 | -0.15 | 0.0201 | 1.03E-13 | -0.217 | 0.0226 | 1.28E-21 |
| Myoglobin | -0.146 | 0.0207 | 1.78E-12 | -0.099 | 0.0233 | 2.33E-05 |
| TIMP1 | -0.143 | 0.0207 | 4.25E-12 | -0.151 | 0.0232 | 9.13E-11 |
| Resistin | -0.268 | 0.0404 | 3.33E-11 | -0.199 | 0.0228 | 3.01E-18 |
| B2M | -0.137 | 0.0208 | 4.00E-11 | -0.144 | 0.0235 | 9.30E-10 |
| EFEMP1 | -0.179 | 0.0337 | 9.95E-08 | -0.182 | 0.0230 | 4.00E-15 |
| FGF23 | -0.103 | 0.0208 | 7.57E-07 | -0.085 | 0.0236 | 3.26E-04 |
| CNTN1 | -0.161 | 0.0408 | 7.84E-05 | -0.128 | 0.0227 | 1.70E-08 |
| Hemopexin | -0.077 | 0.0202 | 0.00014 | -0.095 | 0.0229 | 3.65E-05 |
| PAI1 | 0.094 | 0.03 | 0.00185 | 0.041 | 0.0216 | 0.056 |
| MMP8 | -0.105 | 0.049 | 0.03259 | -0.096 | 0.0234 | 4.29E-05 |

All participants: meta-analysis, fixed-effect (if heterogeneity I^2<0.5) or random-effect (I^2⩾0.5) model to combine summary statistics from discovery and validation sets; Mixed multivariable-adjusted model, adjusted for age, sex, body mass index, smoking, and cohort index, and family structure.

**Supplemental Table 3**. Association results of alcohol intake categories with alcohol related proteins

| Protein | Category | Beta | SE | pvalue | FDR p |
|---|---|---|---|---|---|
| APOA1 | cat1 | 0.199 | 0.0267 | 1.21E-13 | 4.82E-13 |
| APOA1 | cat2 | 0.463 | 0.0416 | 1.69E-28 | 3.38E-27 |
| APOA1 | cat3 | 0.693 | 0.0489 | 8.36E-45 | 5.02E-43 |
| Resistin | cat1 | -0.150 | 0.0267 | 2.04E-08 | 4.08E-08 |
| Resistin | cat2 | -0.340 | 0.0414 | 2.83E-16 | 1.54E-15 |
| Resistin | cat3 | -0.610 | 0.0489 | 2.56E-35 | 7.69E-34 |
| MPO | cat1 | -0.144 | 0.0263 | 4.63E-08 | 8.96E-08 |
| MPO | cat2 | -0.355 | 0.0408 | 3.74E-18 | 3.74E-17 |
| MPO | cat3 | -0.415 | 0.0481 | 8.07E-18 | 6.05E-17 |
| EFEMP1 | cat1 | -0.171 | 0.0270 | 2.45E-10 | 7.35E-10 |
| EFEMP1 | cat2 | -0.254 | 0.0419 | 1.41E-09 | 3.39E-09 |
| EFEMP1 | cat3 | -0.254 | 0.0494 | 2.73E-07 | 4.97E-07 |
| ANGPTL3 | cat1 | -0.162 | 0.0258 | 3.82E-10 | 1.09E-09 |
| ANGPTL3 | cat2 | -0.311 | 0.0401 | 9.73E-15 | 4.49E-14 |
| ANGPTL3 | cat3 | -0.356 | 0.0473 | 6.17E-14 | 2.64E-13 |
| CD56 | cat1 | -0.117 | 0.0265 | 9.58E-06 | 1.34E-05 |
| CD56 | cat2 | -0.279 | 0.0410 | 1.12E-11 | 3.73E-11 |
| CD56 | cat3 | -0.402 | 0.0485 | 1.58E-16 | 9.45E-16 |
| sRAGE | cat1 | -0.089 | 0.0264 | 0.0007 | 0.0009 |
| sRAGE | cat2 | -0.263 | 0.0411 | 1.73E-10 | 5.45E-10 |
| sRAGE | cat3 | -0.518 | 0.0485 | 2.15E-26 | 3.23E-25 |
| FBN | cat1 | -0.163 | 0.0265 | 8.84E-10 | 2.21E-09 |
| FBN | cat2 | -0.346 | 0.0413 | 6.84E-17 | 4.56E-16 |
| FBN | cat3 | -0.243 | 0.0487 | 5.80E-07 | 9.66E-07 |
| Notch1 | cat1 | -0.108 | 0.0273 | 7.66E-05 | 0.0001 |
| Notch1 | cat2 | -0.310 | 0.0425 | 3.08E-13 | 1.15E-12 |
| Notch1 | cat3 | -0.436 | 0.0501 | 4.67E-18 | 4.01E-17 |
| GDF15 | cat1 | -0.240 | 0.0265 | 1.70E-19 | 2.05E-18 |
| GDF15 | cat2 | -0.242 | 0.0411 | 4.00E-09 | 8.89E-09 |
| GDF15 | cat3 | -0.086 | 0.0485 | 0.0754 | 0.0785 |
| Osteocalcin | cat1 | -0.095 | 0.0270 | 0.0004 | 0.0005 |
| Osteocalcin | cat2 | -0.210 | 0.0420 | 6.13E-07 | 9.94E-07 |
| Osteocalcin | cat3 | -0.305 | 0.0494 | 7.48E-10 | 2.04E-09 |
| B2M | cat1 | -0.131 | 0.0274 | 1.75E-06 | 2.57E-06 |
| B2M | cat2 | -0.223 | 0.0425 | 1.57E-07 | 2.94E-07 |
| B2M | cat3 | -0.206 | 0.0502 | 4.26E-05 | 5.81E-05 |
| MMP8 | cat1 | -0.099 | 0.0273 | 0.0003 | 0.0003 |
| MMP8 | cat2 | -0.133 | 0.0424 | 0.0017 | 0.0020 |
| MMP8 | cat3 | -0.159 | 0.0500 | 0.0015 | 0.0017 |
| Cystatin-C | cat1 | -0.139 | 0.0271 | 3.00E-07 | 5.29E-07 |
| Cystatin-C | cat2 | -0.245 | 0.0421 | 6.18E-09 | 1.32E-08 |

| | | | | | |
|---|---|---|---|---|---|
| Cystatin-C | cat3 | -0.248 | 0.0496 | 5.76E-07 | 9.66E-07 |
| TIMP1 | cat1 | -0.161 | 0.0273 | 3.79E-09 | 8.75E-09 |
| TIMP1 | cat2 | -0.209 | 0.0423 | 8.45E-07 | 1.33E-06 |
| TIMP1 | cat3 | -0.244 | 0.0500 | 1.08E-06 | 1.67E-06 |
| Myoglobin | cat1 | -0.049 | 0.0272 | 0.0699 | 0.0749 |
| Myoglobin | cat2 | -0.204 | 0.0423 | 1.36E-06 | 2.05E-06 |
| Myoglobin | cat3 | -0.346 | 0.0498 | 4.01E-12 | 1.42E-11 |
| PAI1 | cat1 | 0.006 | 0.0251 | 0.8092 | 0.8092 |
| PAI1 | cat2 | 0.069 | 0.0390 | 0.0759 | 0.0785 |
| PAI1 | cat3 | 0.282 | 0.0460 | 8.82E-10 | 2.21E-09 |
| FGF23 | cat1 | -0.069 | 0.0275 | 0.0119 | 0.0132 |
| FGF23 | cat2 | -0.144 | 0.0428 | 0.0008 | 0.0009 |
| FGF23 | cat3 | -0.233 | 0.0505 | 3.96E-06 | 5.66E-06 |
| CNTN1 | cat1 | -0.065 | 0.0265 | 0.0136 | 0.0148 |
| CNTN1 | cat2 | -0.233 | 0.0411 | 1.46E-08 | 3.02E-08 |
| CNTN1 | cat3 | -0.382 | 0.0485 | 3.62E-15 | 1.81E-14 |
| Hemopexin | cat1 | -0.085 | 0.0266 | 0.0014 | 0.0016 |
| Hemopexin | cat2 | -0.155 | 0.0415 | 0.0002 | 0.0002 |
| Hemopexin | cat3 | -0.084 | 0.0487 | 0.0841 | 0.0855 |

Linear mixed multivariable-adjusted model, adjusted for age, sex, body mass index, smoking, and cohort index, andfamily structure. SE, standard error; FDR p, false discovery rate adjusted p-value. Cat1: light drinkers (0.1-28 g/day in men and 0.1-14 g/day in women); cat2: at-risk drinkers (28.1-42 g/day in men and 14.1-28 g/day in women); cat3: heavy drinkers (>42 g/day in men and >28 g/day in women); reference group: non-drinkers.

**Supplemental Table 4**. Mendelian randomization of alcohol consumption with alcohol associated proteins

| Protein | nsnp | Beta | Se | pval | FDR |
|---|---|---|---|---|---|
| *APOA1 | 49 | -1.171 | 0.477 | 0.018 | 0.119 |
| Resistin | 49 | 0.149 | 0.271 | 0.583 | 0.995 |
| sRAGE | 49 | 0.018 | 0.287 | 0.949 | 0.995 |
| ANGPTL3 | 49 | -0.837 | 0.292 | 0.004 | 0.059 |
| NOTCH1 | 49 | 0.006 | 0.249 | 0.981 | 0.995 |
| CNTN1 | 49 | 0.035 | 0.282 | 0.900 | 0.995 |
| MPO | 49 | 0.097 | 0.334 | 0.772 | 0.995 |
| CD56 | 49 | 0.457 | 0.362 | 0.206 | 0.687 |
| FBN | 49 | 0.043 | 0.273 | 0.875 | 0.995 |
| Cystatin-C | 49 | 0.217 | 0.288 | 0.450 | 0.995 |
| EFEMP1 | 49 | 0.386 | 0.295 | 0.191 | 0.687 |
| Myoglobin | 49 | 0.317 | 0.320 | 0.322 | 0.877 |
| Osteocalcin | 49 | -0.002 | 0.257 | 0.995 | 0.995 |
| TIMP1 | 49 | 0.066 | 0.273 | 0.809 | 0.995 |
| GDF15 | 49 | -0.042 | 0.289 | 0.883 | 0.995 |
| B2M | 49 | 0.239 | 0.257 | 0.351 | 0.877 |
| FGF23 | 49 | 0.489 | 0.278 | 0.078 | 0.391 |
| PAI1 | 49 | -0.121 | 0.298 | 0.685 | 0.995 |
| MMP8 | 49 | -0.080 | 0.264 | 0.761 | 0.995 |
| Hemopexin | 49 | -0.746 | 0.271 | 0.006 | 0.059 |

FDR: false discovery rate adjusted p-value. * indicates pleiotropy P<0.05.

**Supplemental Table 5**. Gene enrichment analysis of 20 alcohol-associated proteins

| GO biological process complete | # | # | expected | Fold Enrichment | +/- | raw P value | FDR |
|---|---|---|---|---|---|---|---|
| negative regulation of cell adhesion molecule production | 4 | 2 | 0 | > 100 | + | 1.31E-05 | 4.15E-03 |
| post-embryonic eye morphogenesis | 7 | 2 | 0.01 | > 100 | + | 3.13E-05 | 7.54E-03 |
| post-embryonic animal organ morphogenesis | 9 | 2 | 0.01 | > 100 | + | 4.77E-05 | 1.03E-02 |
| modulation of age-related behavioral decline | 9 | 2 | 0.01 | > 100 | + | 4.77E-05 | 1.04E-02 |
| regulation of cell adhesion molecule production | 9 | 2 | 0.01 | > 100 | + | 4.77E-05 | 1.05E-02 |
| negative regulation of extracellular matrix organization | 11 | 2 | 0.01 | > 100 | + | 6.76E-05 | 1.34E-02 |
| phospholipid homeostasis | 13 | 2 | 0.01 | > 100 | + | 9.09E-05 | 1.57E-02 |
| post-embryonic animal morphogenesis | 14 | 2 | 0.01 | > 100 | + | 1.04E-04 | 1.68E-02 |
| cellular response to vitamin D | 15 | 2 | 0.01 | > 100 | + | 1.18E-04 | 1.80E-02 |
| post-embryonic animal organ development | 17 | 2 | 0.02 | > 100 | + | 1.48E-04 | 2.14E-02 |
| negative regulation of lipase activity | 17 | 2 | 0.02 | > 100 | + | 1.48E-04 | 2.15E-02 |
| negative regulation of interleukin-10 production | 19 | 2 | 0.02 | > 100 | + | 1.81E-04 | 2.44E-02 |
| regulation of lipoprotein lipase activity | 22 | 2 | 0.02 | 94.78 | + | 2.38E-04 | 2.98E-02 |
| negative regulation of tissue remodeling | 23 | 2 | 0.02 | 90.66 | + | 2.58E-04 | 3.11E-02 |
| cellular response to vitamin | 24 | 2 | 0.02 | 86.88 | + | 2.80E-04 | 3.25E-02 |
| regulation of heterotypic cell-cell adhesion | 25 | 2 | 0.02 | 83.4 | + | 3.02E-04 | 3.45E-02 |
| positive regulation of lipid catabolic process | 27 | 2 | 0.03 | 77.23 | + | 3.49E-04 | 3.82E-02 |
| negative regulation of biomineral tissue development | 28 | 2 | 0.03 | 74.47 | + | 3.73E-04 | 3.86E-02 |
| positive regulation of mononuclear cell migration | 28 | 2 | 0.03 | 74.47 | + | 3.73E-04 | 3.88E-02 |
| negative regulation of biomineralization | 28 | 2 | 0.03 | 74.47 | + | 3.73E-04 | 3.91E-02 |
| response to vitamin D | 30 | 2 | 0.03 | 69.5 | + | 4.25E-04 | 4.36E-02 |
| protein-lipid complex remodeling | 31 | 2 | 0.03 | 67.26 | + | 4.52E-04 | 4.50E-02 |
| plasma lipoprotein particle remodeling | 31 | 2 | 0.03 | 67.26 | + | 4.52E-04 | 4.52E-02 |
| protein-containing complex remodeling | 32 | 2 | 0.03 | 65.16 | + | 4.80E-04 | 4.69E-02 |

**Supplemental Table 6**. Alcohol related proteins associations with cardiovascular outcomes from Ho's paper[9]

| Trait | Biomarker | HR | 95% LCL | 95% UCL | *P* value |
|---|---|---|---|---|---|
| Atheroslerotic CVD (N=392) | GDF15 | 1.38 | 1.20 | 1.58 | 3.6E-06 |
| | TIMP1 | 1.32 | 1.17 | 1.48 | 5.5E-06 |
| | B2M | 1.24 | 1.10 | 1.40 | 3.6E-04 |
| | Cystatin-C | 1.21 | 1.08 | 1.36 | 1.1E-03 |
| Heart failure (N=226) | GDF15 | 2.08 | 1.72 | 2.53 | 9.1E-14 |
| | B2M | 1.47 | 1.24 | 1.73 | 4.5E-06 |
| | Cystatin-C | 1.43 | 1.22 | 1.67 | 6.5E-06 |
| | TIMP1 | 1.39 | 1.18 | 1.63 | 6.0E-05 |
| | MPO | 1.30 | 1.13 | 1.48 | 1.4E-04 |
| | EFEMP1 | 1.34 | 1.14 | 1.57 | 3.2E-04 |
| | Resistin | 1.25 | 1.09 | 1.43 | 1.1E-03 |
| All-cause mortality (N=755) | GDF15 | 1.96 | 1.76 | 2.17 | 1.2E-35 |
| | B2M | 1.39 | 1.27 | 1.52 | 1.0E-12 |
| | TIMP1 | 1.36 | 1.24 | 1.48 | 1.0E-11 |
| | EFEMP1 | 1.28 | 1.17 | 1.40 | 2.6E-08 |
| | Cystatin-C | 1.23 | 1.13 | 1.34 | 1.9E-06 |
| | FGF23 | 1.19 | 1.10 | 1.28 | 8.0E-06 |
| | MMP8 | 1.17 | 1.08 | 1.26 | 4.1E-05 |
| | MPO | 1.12 | 1.04 | 1.21 | 1.7E-03 |
| | Resistin | 1.12 | 1.04 | 1.20 | 2.8E-03 |
| | CNTN1 | 0.90 | 0.83 | 0.97 | 7.9E-03 |
| CVD death (N=167) | GDF15 | 1.96 | 1.56 | 2.46 | 6.9E-09 |
| | B2M | 1.72 | 1.42 | 2.09 | 3.0E-08 |
| | EFEMP1 | 1.60 | 1.33 | 1.94 | 9.5E-07 |
| | Cystatin-C | 1.55 | 1.29 | 1.87 | 4.1E-06 |
| | TIMP1 | 1.40 | 1.16 | 1.69 | 5.0E-04 |
| | Resistin | 1.28 | 1.09 | 1.49 | 2.2E-03 |
| | FGF23 | 1.28 | 1.09 | 1.50 | 2.6E-03 |
| | FBN | 1.25 | 1.06 | 1.48 | 9.4E-03 |

Multivariable-adjusted model, adjusted for age, sex, systolic blood pressure, hypertension treatment, diabetes mellitus, body mass index, smoking, total and HDL cholesterol, and history of atrial fibrillation. In addition, heart failure and mortality analyses were adjusted for prevalent myocardial infarction. CVD indicates cardiovascular disease; HDL, high-density lipoprotein; HR, hazards ratio per 1-SD change in rank normalized data; LCL, lower 95% confidence interval; UCL, upper 95% confidence interval

**Supplemental Figure 1a**. Three-way association of alcohol intake, alcohol related proteins, and CVD risk factors as expected

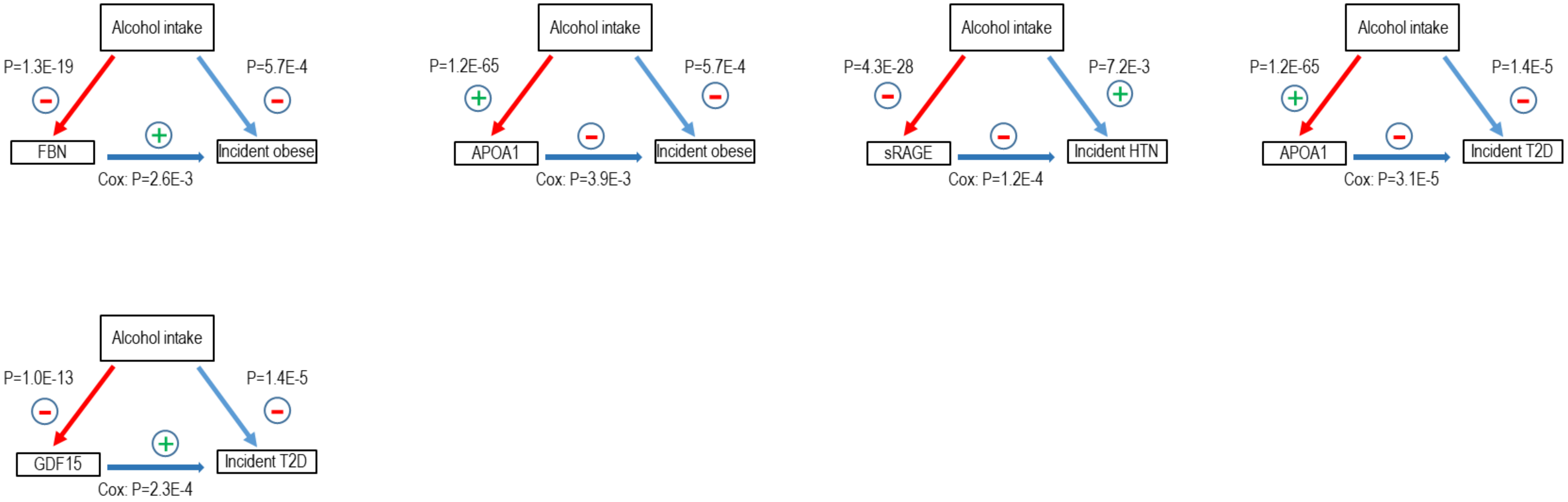

**Supplemental Figure 1b**. Three-way association of alcohol intake, alcohol related proteins, and CVD risk factors not as expected

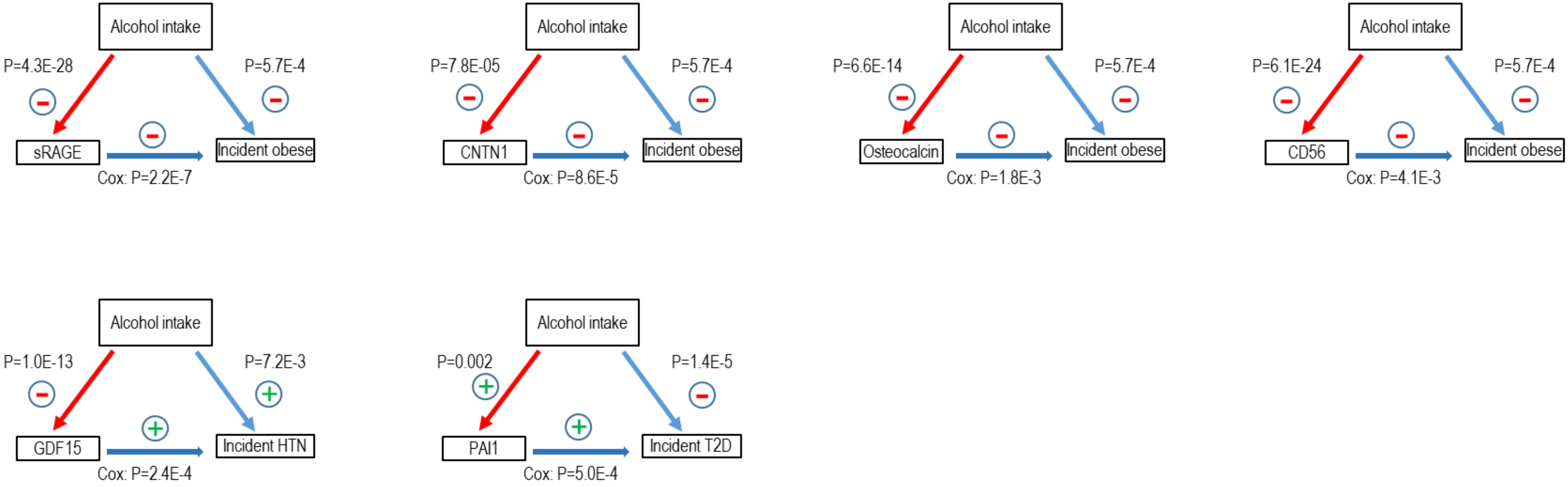
